# Pleiotropic function of *Dlx5/6* in coordinating the development of mammalian vocal and auditory organs

**DOI:** 10.1101/2024.12.13.628362

**Authors:** Frida Sánchez-Garrido, Victoria Bouzerand, Markéta Kaiser, Chloé Chaumeton, Anastasia Fontaine, Tomáš Zikmund, Jozef Kaiser, Églantine Heude

**Author notes:** To whom correspondence should be addressed: Dr. Eglantine Heude, Institut de Génomique Fonctionnelle de Lyon, CNRS UMR 5242, 32-34 avenue Tony Garnier 69007 Lyon.

## Abstract

Acoustic communication, essential for mammalian social interactions, involves both effector (vocal) and receptor (auditory) organs that present a wide variety of morphologies between species. The molecular mechanisms supporting the harmonious diversification of effector and receptor systems along with the evolution of species-specific acoustic communication are still poorly understood. It is conceivable that common genetic pathways determine the parallel morphogenesis of vocal and auditory systems. Here, we investigated this question by generating mutant mice with targeted invalidation of *Dlx5/6* genes in the otic placode and cephalic neural crest cells, contributing to the formation of the ear and vocal tract components. We show that *Dlx5/6* inactivation provokes simultaneous defects of the outer, middle and inner ear and of the jaw, pharynx and larynx musculoskeletal systems. Our findings support the notion that *Dlx5/6* genes played a pleiotropic role in the co-adaptation of vocal and auditory systems in mammals.

## Introduction

Acoustic communication is remarkably different between land vertebrate (tetrapod) species going from ultrasonic signals in mice and bats to speech in humans. It invariably involves an effector organ, the vocal tract which produces sounds, and a receptor organ, the auditory system, that receives sounds^1–5^. Acoustic communication is critical to share information and for social interactions, such as sexual selection, maternal care or species recognition. The need for complementarity between sound emission and reception suggests a coordinated evolution and development of organs supporting these functions.

The vocalization process engages the coordination of a set of cranio-cervical organs including the jaws, the tongue, the pharynx and the larynx^4^. In parallel, sounds are treated through the outer (pinna), middle (ossicles) and inner (cochlea) components of the ear. In the mammalian embryo, the vocal tract originates from the pharyngeal arches (PAs)^6–10^. During early embryogenesis, these transient structures are colonised by multiple cell populations, including the cephalic neural crest cells (CNCC), the cardiopharyngeal mesoderm and mesodermal cells derived from anterior somites. CNCC gives rise to the jaws, the hyoid bone and the thyroid cartilage of the larynx, and to associated connective tissues and tendons^7,10–12^. Besides that, the auditory system derives from the otic vesicle and part of the PAs. While the inner ear derives from the ectodermal otic placode, the outer ear, the ossicles of the middle ear (stapes, incus malleus) and the tympanic ring derive from PA1-2 CNCC^13,14^.

In pharyngeal arches, CNCC patterning is regulated by the *Dlx* gene family, homologous to the drosophila *distal-less* (*dll*) gene, coding for homeodomain transcription factors. In mammals, the *Dlx* family consists of three tandems of genes (*Dlx1/2, Dlx3/4* and *Dlx5/6),* with genes from the same pair showing similar expression profile and redundant functions^15–17^. During embryonic development, *Dlx5* and *Dlx6* are sequentially expressed at the lateral plate border of the neural tube, within PAs in migrating and post-migrating CNCC, in the apical ectodermal ridge of the limb bud, and later during skeletal differentiation^15,17–19^. Studies have demonstrated that *Dlx5/6* expression is required to specify mandibular identity as their constitutive inactivation leads to a transformation of the lower jaw into an upper jaw^15,19,20^. Constitutive and conditional inactivation of *Dlx5/6* in CNCC also induce defects of tongue musculature and an absence of masticatory muscles of mesodermal origin^9,15^. *Dlx5/6* genes have thus an instructive role in coordinating the formation of the jaw musculoskeletal system. *Dlx5/6^-/-^* mice also present neural tube defects leading to exencephaly and to severe cranial malformations^15,20,21^. In addition, malformations of the vestibular inner ear, the tympanic ring, the middle ear ossicles, the pinna, and of the hyoid and thyroid cartilages have been occasionally described in *Dlx5/6* mutants, but the overall cranio-cervical defects have never been fully addressed^9,15,18,19,22,23^.

In 1910, Ludwig Plate described the concept of “pleiotropy” where a single genetic locus could affect multiple phenotypic traits^24^. Since then, pleiotropy has been a central concept in the fields of medical genetics, evolutionary and developmental biology^25,26^. Given the key role of *Dlx5/6* genes in the specification of lower jaw identity, of some components of the vocal tract and of the auditory apparatus, here we investigated the potential pleiotropic role of the genes in the formation of the vocal and auditory apparatus at the basis of acoustic communication in mammals.

To do so, we generated conditional mutant mice with targeted invalidation of *Dlx5/6* genes in post migratory CNCC and in the otic vesicle using the *Sox10-Cre* driver^27,28^. Using histological approaches and X-ray computed microtomography (micro-CT), we show that *Dlx5/6* conditional inactivation induces defects of the musculoskeletal system of all organs composing the vocal tract, and of the external, middle and inner ear. By investigating morphogenetic events at the origin of malformations, our results reveal that *Dlx5/6* invalidation affects the early patterning of the otic vesicle and of CNCC derivatives within the PAs, without affecting their cellular fate. We also show that Dlx5/6 transcription factors regulate the expression of actors of the BMP signalling pathway in both PAs and otic vesicles. Altogether, our data unveil the pleiotropic role of *Dlx5/6* in the formation of all components of the acoustic vocal and auditory complex. We propose that a common Sox10-Dlx5/6-BMP axis contributed to the co-adaptation of the effector and receptor organs during acoustic communication diversification in land vertebrates.

## Results

### Dlx5/6 invalidation affects the entire vocal tract and auditory systems

To induce targeted invalidation of *Dlx5/6* in CNCC and otic vesicle (*Dlx5/6* cKO), we crossed *Sox10^Cre/+^* deleter mice with the previously-described *Dlx5/6^flox/flox^* mouse line^9,29^, in which both the homeodomain-encoding region (exon 2) of *Dlx5* and *Dlx6* are flanked by *loxP* sequences. *Sox10* and *Dlx5/6* genes are both expressed in post migrating CNCC populations, in the otic placodes and their derivatives (Supp. Fig. 1) ^15,18,27,28^.

The macroscopic phenotypic analysis before birth revealed that *Dlx5/6* cKO foetuses presented craniofacial malformations similar to those observed in previous *Dlx5/6* invalidated mouse models^9,15,18,19^, including a transformed lower jaw exhibiting ectopic vibrissae and hypoplastic external ear (Fig. 1A-A’).

**Figure 1.**
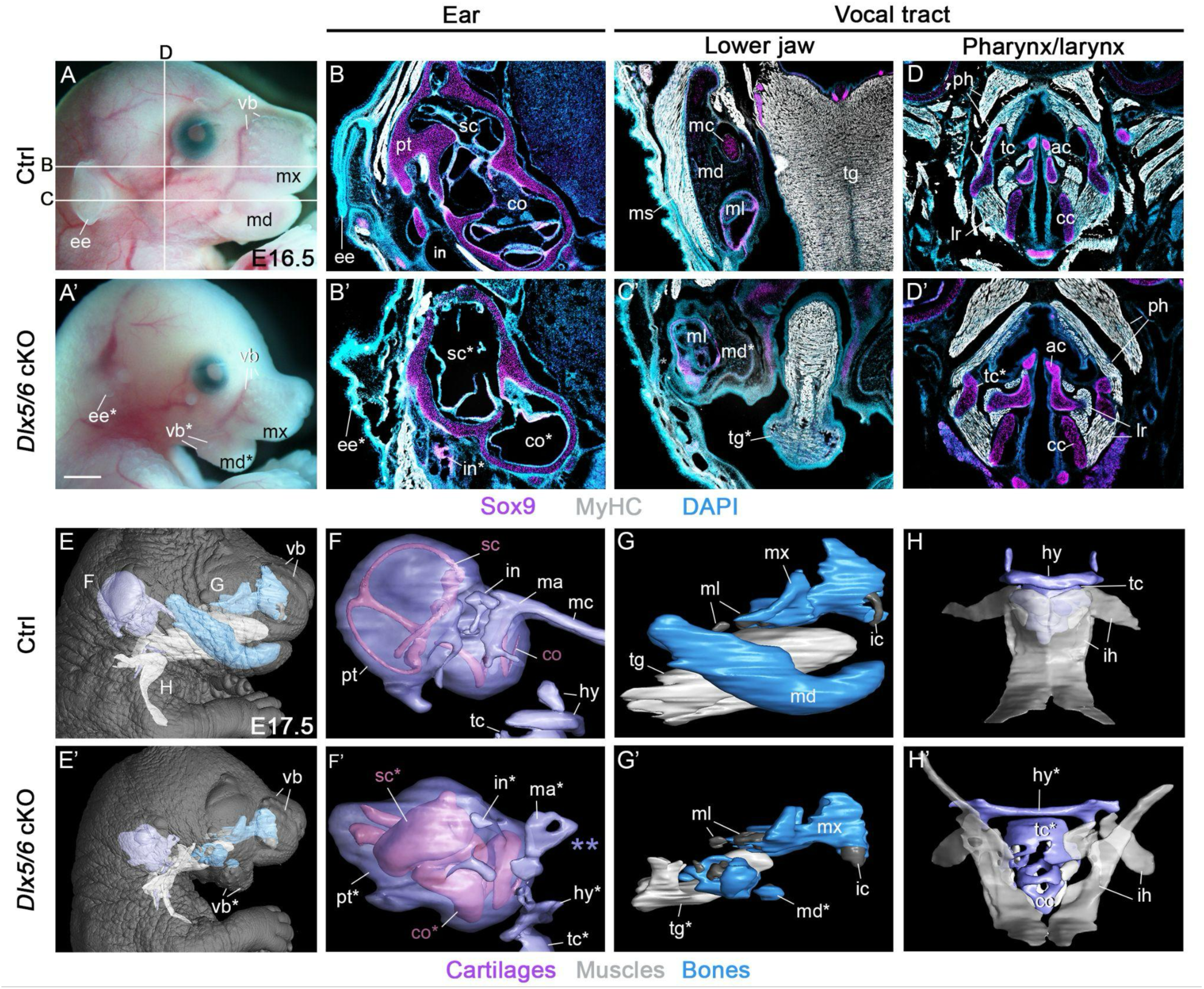
*Dlx5/6* genes are required for the proper formation of organs composing both vocal and auditory systems. (A-A’) Macroscopic view of control and *Dlx5/6* cKO mutant phenotypes at E16.5. In mutants, the lower jaw is defective with ectopic vibrissae (vb*) and the external ear (pinna) shows severe hypoplasia. (B-D’) Immunofluorescence stainings for the cartilage marker Sox9 and muscle marker MyHC on coronal and frontal sections at the levels indicated in A. (B-B’) The *Dlx5/6* mutant presents malformations of external, middle and inner ear components including reduced pinna, defective incus and semicircular canals and enlarged cochlear duct. (C-D’) The *Dlx5/6* mutant presents a transformed lower jaw characterized by an absence of Meckel cartilage and of associated masticatory musculature. The tongue is severely reduced with misorganized myofibers. The thyroid cartilage also seems reduced compared to control (n=6). (E-H’) 3D reconstructions of the different structures composing the vocal tract and auditory systems in control and *Dlx5/6* mutant conditions at E17.5 (n=2). (F-F’) The Meckel cartilage is missing (purple double asterisk in F’), the middle ear ossicles (incus and malleus) are defective and the vestibular and cochlear membranous labyrinths are swollen in mutants compared to controls. (G-H’) The lower jaw transformation in mutants is associated with tongue hypoplasia and to a mispatterning of extrinsic laryngeal (infrahyoid) muscles. The thyroid and hyoid cartilages also show defects of formation in the mutant context. All structures presenting a malformation in the *Dlx5/6* mutant context are indicated with an asterisk. Absent or reduced structures are noted with a double asterisk. Abbreviations: mx, maxillary; md, mandible; vb, vibrissae; ee, external ear; pt, petrous part of the temporal bone; sc, semicircular canals; co, cochlea; ms, masseter muscle; mc, Meckel cartilage; ml, molar; ic, incisor; tg, tongue; ph, pharyngeal muscles; lr, laryngeal muscles; tc, thyroid cartilage; hy, hyoid cartilage; ac, arytenoid cartilage; cc, cricoid cartilage; in, incus; ma, malleus. Scale bar in A’ for A-A’ 1000 µm, for B-D’ 200 µm.

First, we performed immunofluorescence stainings for MyHC and Sox9 on sections to investigate respectively the muscular and skeletal phenotypes of control and mutant foetuses at E16.5. Regarding the auditory system, Sox9 immunostainings revealed an hypoplasia of the middle ear ossicles of CNCC origin^13^ (Fig. 1B-B’). The inner ear showed defects of the membranous and bony labyrinths of the semicircular canals and cochlea (Fig. 1B-B’). At the level of the vocal tract, we observed that the transformed lower jaw was associated with tongue hypoplasia and masticatory masseter muscle aplasia (Fig. 1C-C’), as previously described after *Dlx5/6* inactivation^9,15^. The hyoid and the laryngeal thyroid cartilages derived from CNCC^7,12^ seemed also impacted but the extent of malformation was difficult to access on sections.

To push forward the phenotypic analysis, we performed micro-CT scanning of mutant and control foetuses at E17.5 to access the complex layout of the craniocervical structures and to reconstruct in 3D the different components of the vocal tract and auditory system (Fig. 1E-H’). This approach clearly revealed the extent of inner ear malformations including swollen membranous labyrinths with indiscernible semicircular canals and wider cochlear duct (Fig. 1F-F’). Moreover, we observed the presence of two small mispatterned skeletal structures at the site of the incus and malleus ossicles associated with an absence of Meckel cartilage in mutants (Fig. 1F-F’). The 3D reconstruction of the jaw musculoskeletal elements highlighted the severe malformations of the lower jaw (mandible) and of the tongue musculature (fig 1, G-G’). The hyoid and thyroid cartilages were severely affected and extrinsic laryngeal (infrahyoid) muscles showed patterning defects (Fig. 1H-H’). Pharyngeal and intrinsic laryngeal muscles presented morphology differences compared to controls adapted to the defective hyoid and thyroid cartilages to which they connect. In contrast, the cricoid and arytenoid cartilages of mesodermal origin^7,12^ appeared unaffected by *Dlx5/6* mutation (Fig. 1D-D’, Supp. Data 2-3).

### Dlx5/6 genes orchestrate early CNCC and otic vesicle patterning

We then used genetic lineage tracing approaches to analyze the morphogenetic events at the basis of the musculoskeletal malformations in *Dlx5/6* mutants. We first examined the fate of CNCC by *Sox10* genetic lineage tracing using the *Rosa^lolx-stop-lox-lacZ^* (*Rosa^lacZ/+^*)^30^ reporter analysis at E11.5 in control and mutant conditions (Fig. 2A-B). β-galactosidase (β-gal) positive cells corresponding to placode derivatives, were detected in the tympanic ring and the endolymphatic sac of the otic vesicle in controls (Fig. 2A-A’). We also observed β-gal-positive cells corresponding to CNCC derivatives within the mesenchyme of the maxillary and mandibular parts of the PA1, the hyoid arch (PA2) and more posterior arches (PA3-6) (Fig. 2A”), which later give rise to the skeletal components of the jaws, the ossicles and the hyoid and thyroid cartilages. In *Dlx5/6* cKO mutants, both compartments showed organization defects of β-gal-positive cells. In the developing ear, the endolymphatic sac was missing and the tympanic ring was mis-patterned, whereas β-gal-positive CNCC were scattered and disorganised within the PA1-6 (Fig. 2B-B’’). Remarkably, we did not observe any other defects along the body axis, the targeted *Dlx5/6* mutation affecting specifically vocal tract and auditory precursors.

**Fig. 2.**
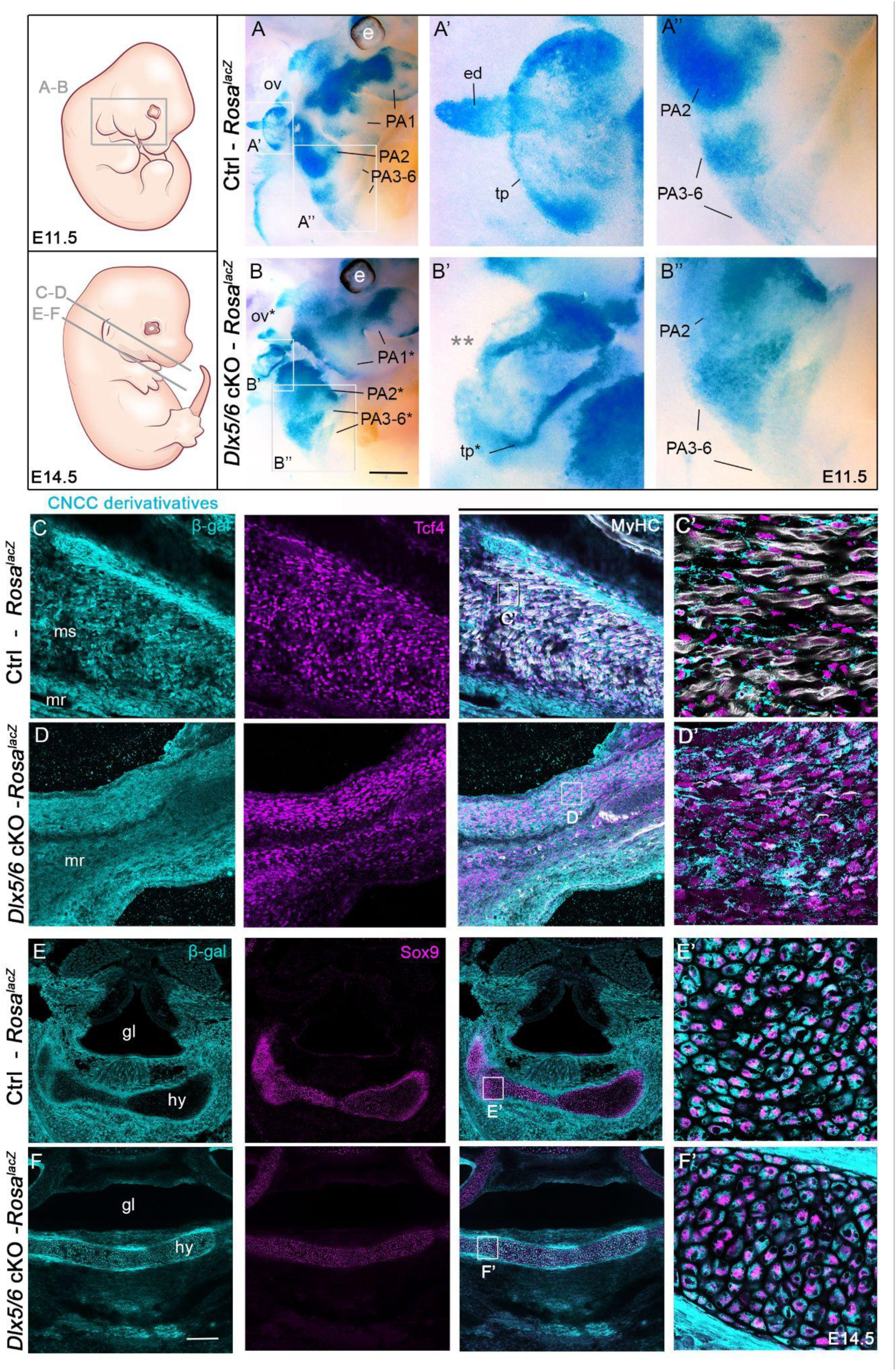
*Dlx5/6* orchestrate the patterning of CNCC in the PAs and of the otic vesicle without affecting cell fate. (A and B) *Sox10* genetic lineage tracing of CNCC and otic placode derivatives using the Rosa-lacZ reporter in control and *Dlx5/6* mutant embryos at E11.5. In mutants, the otic vesicle shows a lack of endolymphatic sac (grey double asterisk in B’) and defect of the tympanic ring compared to controls (A’-B’). β-gal-positive CNCC compose the mesenchymal population of the PAs in controls that show aberrant patterning in mutants (A”-B”). (C-F’) Immunofluorescence stainings on sections for β-galactosidase labelling CCNC derivatives, and for Tcf4, MyHC and Sox9 that mark connective tissue fibroblasts, muscles and cartilages respectively in control and mutant foetuses at E14.5. Tcf4-positive fibroblasts of CNCC origin are detected between masticatory myofibers associated with the lower jaw in controls (C-C’). β-gal/Tcf4-positive fibroblasts are maintained in the masticatory region of mutants despite the absence of differentiated musculature (D-D’). At the level of the glottis and hyoid cartilage, we observe differences in the distribution of CNCC derivatives that still keep their cartilaginous identity (E-F’) (n=3). All structures presenting a malformation are marked with an asterisk. Absent or reduced structures are noted with a double asterisk. Abbreviations: ed, endolymphatic sac; tp, tympanic ring; ms, masseter muscle; mr, masticatory region; gl, glottis; hy, hyoid cartilage; ov, otic vesicle; PA, pharyngeal arch. Scale bar in B for A-B 400 µm, for A’-B’ 150 µm, for A”-B” 200 µm, in F for C-F 200 µm, for C’-F’ 20 µm.

We then followed the fate of β-gal derivatives during musculoskeletal differentiation at E14.5. We performed immunofluorescence stainings on sections for anti-β-gal and for markers of cartilaginous, fibroblastic or tendinous derivatives (Sox9, Tcf4 and Tnc respectively) of placodal and CNCC populations. In controls, CNCC-derived β-gal cells corresponded to Tcf4-positive fibroblasts forming the muscle connective tissue along the myofibers of the masticatory masseter and tongue muscles (Fig. 2C-C’, Supp. Fig. 2A)^7^. In mutants, the β-gal/Tcf4-positive muscle connective tissue fibroblasts are present, even in the absence of masseter musculature (Fig. 2D-D’, Supp. Fig. 2B). The β-gal/Tcf4-positive populations of the external ear were also preserved in the hypoplastic pinna of mutants that however showed defects in organization compared to controls (Supp. Fig. 2C-D). The β-gal-positive CNCC-derived population showed aberrant patterning at the level of the glottis, the hyoid and thyroid cartilages but kept their cartilaginous and tendinous identity as observed by Sox9, Tnc and β-gal colocalization in the structures of both controls and mutants (Fig. 2E-F’, Supp. Fig. 2E-H).

We then investigated whether such alterations in *Dlx5/6* expression could also affect the innervation of vocal tract and auditory components. We analyzed the neuromuscular system of control and mutant embryos by *in toto* immunofluorescence stainings at E11.5 targeting the neurofilament (NF) and Desmin proteins that mark respectively the developing peripheral nervous and muscular systems (Fig. 3). In controls, the maxillary and mandibular branches of the trigeminal ganglion (cranial nerve CN V) project into the upper and lower jaw buds to innervate notably the sensory vibrissae and the masticatory muscle precursor (Fig. 3A-A’). We also observed the facial nerve (CN VII) that innervates the facial muscles and its *chorda tympani* branch that later join the lingual nerve (CN V) in the mandibular arch^31^. More dorsally, the vestibulocochlear nerve (CN VIII) innervates the inner ear, and posteriorly the glossopharyngeal, the vagus and accessory nerves (CN IX, X, XI) project to innervate the tongue, throat and laryngeal components (Fig. 3A-A’, Supp. Video 1) ^32^. In *Dlx5/6* cKO, the mandibular branch of the trigeminal ganglion shows increased distal arborization characteristic of the maxillary branch to innervate the ectopic vibrissae observed on the transformed lower jaw (Fig. 1A’, E’, Fig. 3B). The vestibulocochlear nerve (CN VIII) shows defect of projection on the otic vesicle in the mutants compared to controls (Fig. 3B-B’). Moreover, one branch of the *chorda tympani* that innervates the distal part of tongue later during development was missing. However, we did not detect an obvious difference in the hypoglossal nerve (CN XII) while the hypoglossal cord forming the tongue was already reduced. In contrast, the masticatory muscle precursor was present at this stage as previously described in constitutive *Dlx5/6* mutants, the defect of masticatory muscle differentiation occurring later during development in mutants^15^ (Fig. 2). We did not notice differences in the configuration of the glossopharyngeal, the vagus and accessory nerves (CN IX, X, XI) between controls and mutants (Fig. 3B’-D, Supp. Video 2).

**Fig. 3.**
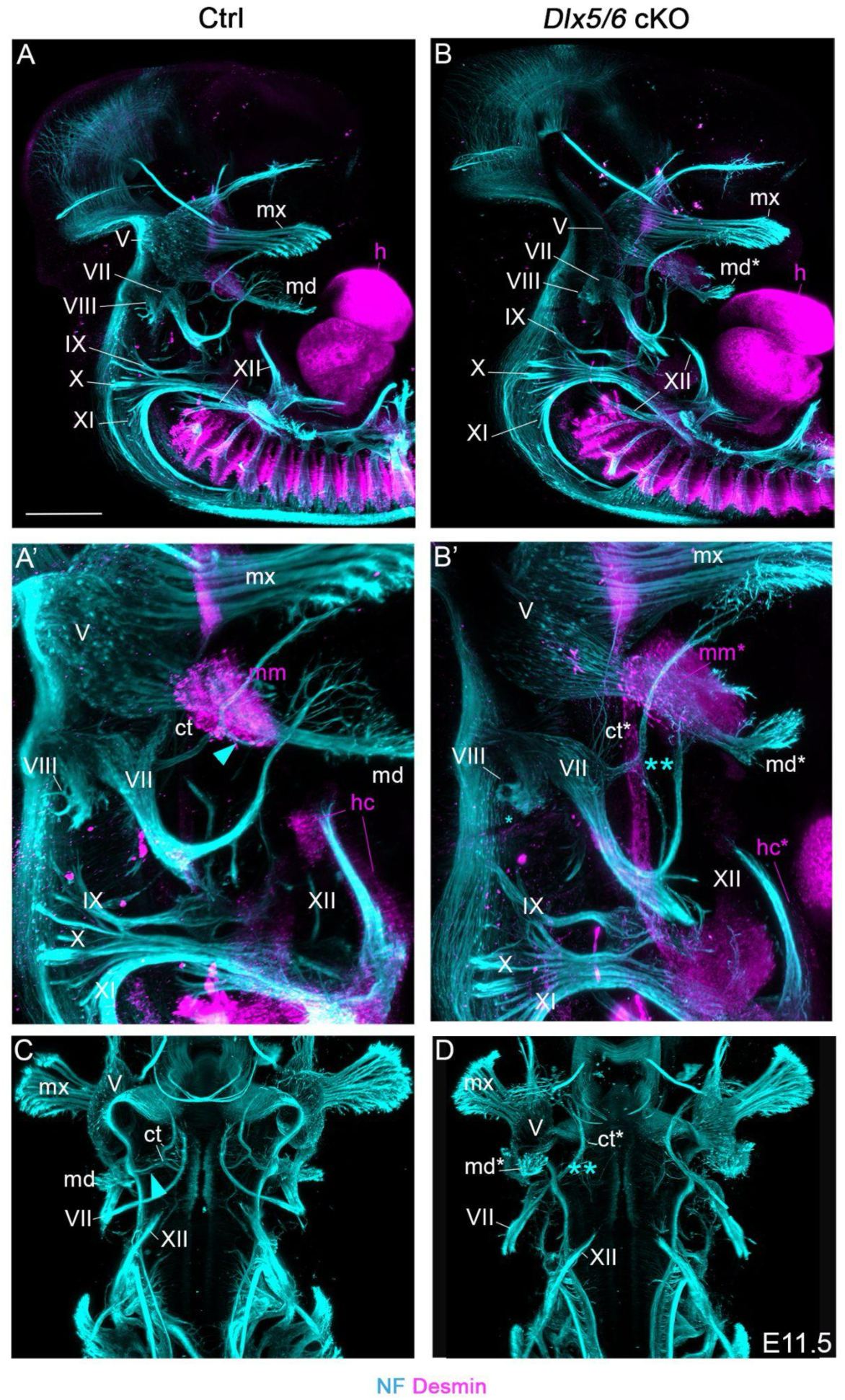
*Dlx5/6* invalidation induces innervation defects of some vocal and auditory components. (A-D) *In toto* immunofluorescence staining of control and mutant embryos at the E11.5 for the neurofilament (NF) and Desmin that label the peripheral nervous system and muscle precursors respectively. Lateral (A-B’) and ventral (C-D) views with magnification at the level of vocal tract and auditory precursors (A’-B’). In mutants, the mandibular branch of the trigeminal nerve (V) presents characteristics of the maxillary branch with increased distal arborization to connect ectopic vibrissae. The vestibulocochlear nerve (VIII) shows defective projection toward the otic vesicle. The *chorda tympani* branch of the facial nerve (VII) (blue arrowheads in controls A’, C) misses the neuronal projection that later innervates the tongue (blue double asterisk) in mutants B’, D). Note that the masticatory muscle precursor is present at E11.5 while the hypoglossal cord giving rise to tongue muscles is already reduced in mutants (n=3). All structures presenting a mutant phenotype are noted with an asterisk. Absent or reduced structures are noted with a double asterisk. Abbreviations: mx, maxillary branch of the trigeminal ganglion; md, mandibular branch of the trigeminal ganglion; h, heart; mm, masticatory muscle precursor; ct, *chorda tympani*; hc, hypoglossal cord. Scale bar in A for A-B, C-D 300 µm, for A’-B’ 200 µm.

Altogether, our results show that *Dlx5/6* expression orchestrates synergistically the proper patterning and innervation of CNCC and otic derivatives without affecting their cell fate during the development of vocal tract and auditory components.

### Dlx5/6 regulate BMP signaling in both pharyngeal arches and otic vesicle

It was previously reported that several actors of the BMP signaling pathway are downstream targets of Dlx5 in the otic vesicle^33^. Some BMP signaling genes were also shown to be expressed during pharyngeal arch patterning and affected by *Dlx5/6* constitutive inactivation^34^. We thus wondered if Dlx5/6 transcription factors may act as regulators of the BMP signaling pathway in both vocal tract and auditory precursors.

We selected key genes, *Bmper, Msx1* and *Hand2*, and analyzed their expression by *in situ* hybridization during early development at E10.5. Bmper (BMP-binding endothelial cell precursor-derived regulator) is a secreted protein present in migrating CNCC and the mesodermal core of PAs^34–37^. It was reported that *Bmper-*inactivated mice showed malformations of the thyroid cartilage^36^. Msx1 and Hand2 are downstream components of the BMP signaling cascade, they are expressed in CNCC and are regulated by *Dlx5/6* in pharyngeal arches^34,38,39^. *Msx1* and *Hand2* deficient mice show malformations of the middle ear, the hyoid and thyroid cartilages^40,41^.

All the genes analyzed were expressed in both otic vesicle and pharyngeal arches at E10.5 in control embryos (Fig. 4). *Bmper* was expressed in the tympanic ring of the otic vesicle and in the mesodermal core of the PA1-2 (Fig. 4A). Expression was also noticed in the distal part of PA1-2 and in PA3-6. In *Dlx5/6* mutant embryos, expression in the distal PA1-2 and otic vesicles was reduced and undetectable in the PA1-2 mesodermal cores and posterior arches (PA3-6) (Fig. 4B).

**Fig. 4.**
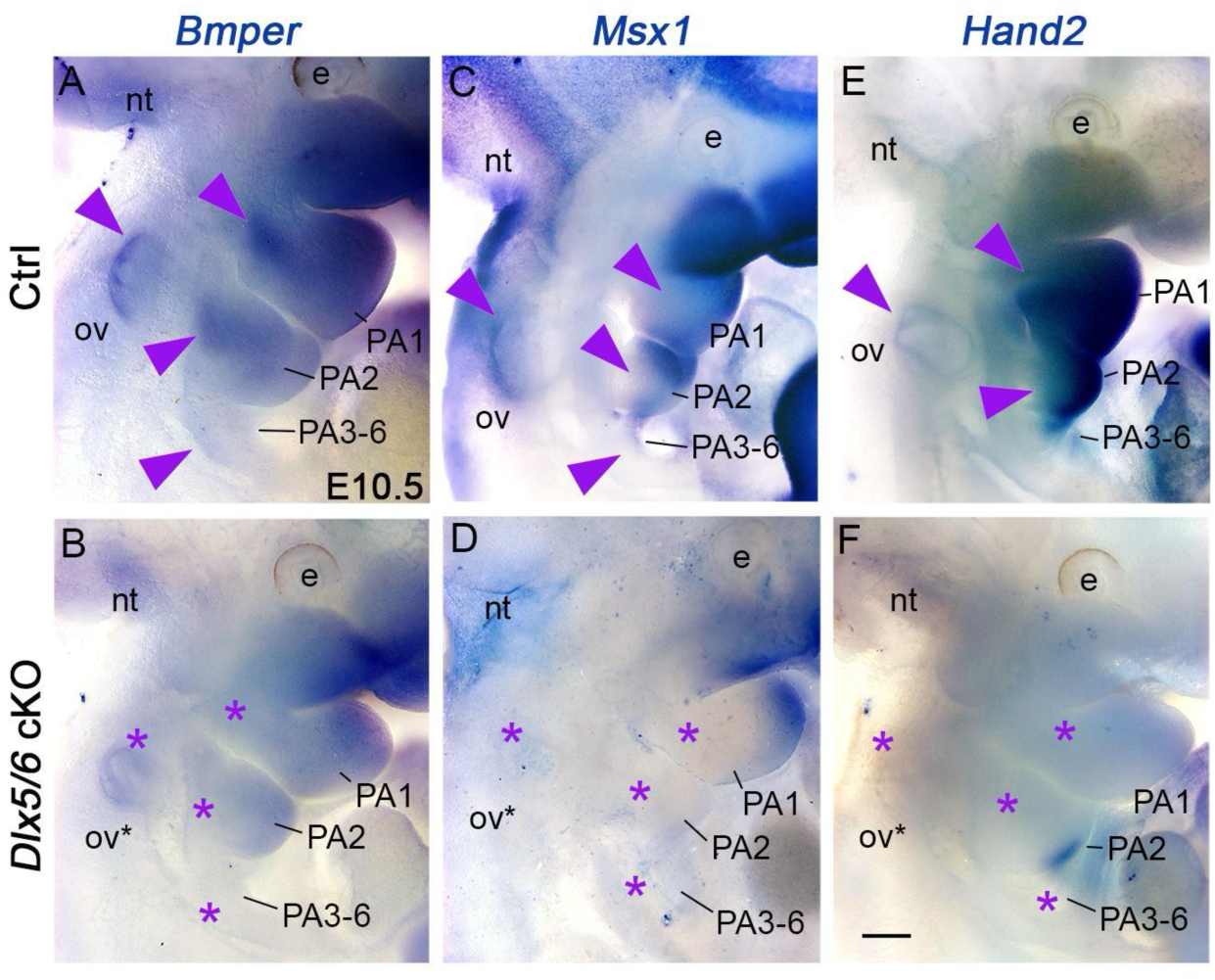
Dlx5/6 transcription factors regulate BMP signaling in both pharyngeal arches and otic vesicle. Whole mount *in situ* hybridization for *Bmper*, *Msx1* and *Hand2* in E10.5 control and mutant embryos (A-F). *Bmper* expression is observed in the mesodermal core and the distal part of PA1-2 and in the tympanic ring of controls (A). In mutants, *Bmper* expression is lost in the mesodermal cores and expression is reduced in the distal PAs and otic vesicle (B). In controls, *Msx1* and *Hand2* are expressed in all PAs and otic vesicles (arrowheads in C, E). In mutants, expression is undetectable in the otic vesicle and reduced or undetectable in PAs (n=4 each condition) (D, F). All structures presenting undetectable or reduced gene expression in the mutant are marked with an asterisk. Abbreviation: nt, neural tube; e, eye; PA, pharyngeal arch; OV, otic vesicle. Scale bar in F for A-F 200 µm.

In the control, *Msx1* was expressed in the distal cap of PA1-2, in posterior PAs, as well as the otic vesicle (Fig. 4C). In our mutants, *Msx1* expression was lost in PA2-6 and otic vesicles, and reduced in PA1 (Fig. 4D). *Hand2* was strongly expressed in all pharyngeal arches and expression was detected in the otic vesicle in controls (Fig. 4E). The *Dlx5/6* mutant embryos showed a drastic loss of expression in both compartments (Fig. 4F). Our analysis indicates that *Dlx5/6* genes directly regulate the expression of actors of the BMP signaling pathway early during the development of both vocal tract and auditory precursors.

## Discussion

### A common Sox10-Dlx5/6-BMP axis coordinates auditory and vocal tract development

In this study, we show that Dlx5/6 transcription factors control the early patterning and innervation of pharyngeal arch and otic derivatives by regulating the BMP signaling pathway in both compartments.

The BMP signaling pathway is implicated in many developmental processes including cellular differentiation, growth and apoptosis^42^. It has been previously shown that *Dlx* genes and BMP signaling interact for proper craniofacial development and notably for the formation of the jaw and inner ear^22,43^. In our study, we selected and analyzed actors of the BMP signaling pathway that were shown to be direct downstream targets of *Dlx5* in the otic placode^33^ and that we noticed being also expressed in both PAs and otic vesicles. We observed that the gene coding for the secreted protein Bmper (Cv2) was expressed in distal CNCC and within the mesodermal cores of PA1-2 in controls. In our mutants, *Bmper* expression was reduced in CNCC and not detectable in the mesodermal cores in which *Dlx5/6* is not expressed, indicating indirect modulation of *Bmper* expression by *Dlx5/6* in the latter compartment. We previously demonstrated that *Dlx5/6* expression in CNCC is necessary to instruct the adjacent mesoderm for the differentiation of masticatory muscles^9,15^, but the molecular actors involved in CNCC-mesoderm interaction remained elusive. We hypothesize that the secreted Bmper, expressed in both CNCC and mesodermal populations and regulated by Dlx5/6, may handle such a role. *Bmper* invalidated mice show defects of laryngeal cartilages and hypoplastic skull vault, however the head muscular phenotype had not been investigated^36^. Further functional analyses in the mouse embryos are still required to test if Bmper could be a mediator between Dlx5/6 in CNCC and the adjacent mesoderm to regulate head muscle differentiation.

We have also shown that inactivation of *Dlx5/6* severely impacts the expression of Hand2 and Msx1 in both otic vesicles and PAs. These transcription factors are known to act downstream of the BMP signaling and it has been proposed that *Dlx5/6* regulate their expression to direct lower jaw patterning^34,39^. Moreover, Chip-seq data have identified them as direct targets of Dlx5 in the inner ear precursor^33^. In contrast to *Wnt1* that is a specific and early marker of neural crest derivatives, the *Sox10* lineage also includes the otic vesicle^28^. In chick, a common enhancer element regulates *Sox10* expression in the otic placode and CNCC^27^. The data thus indicate that a common Sox10-Dlx5/6-BMP signaling axis coordinates the morphogenesis of both CNCC and otic placode derivatives for the proper formation of the vocal and auditory complex (Fig. 5).

**Fig. 5.**
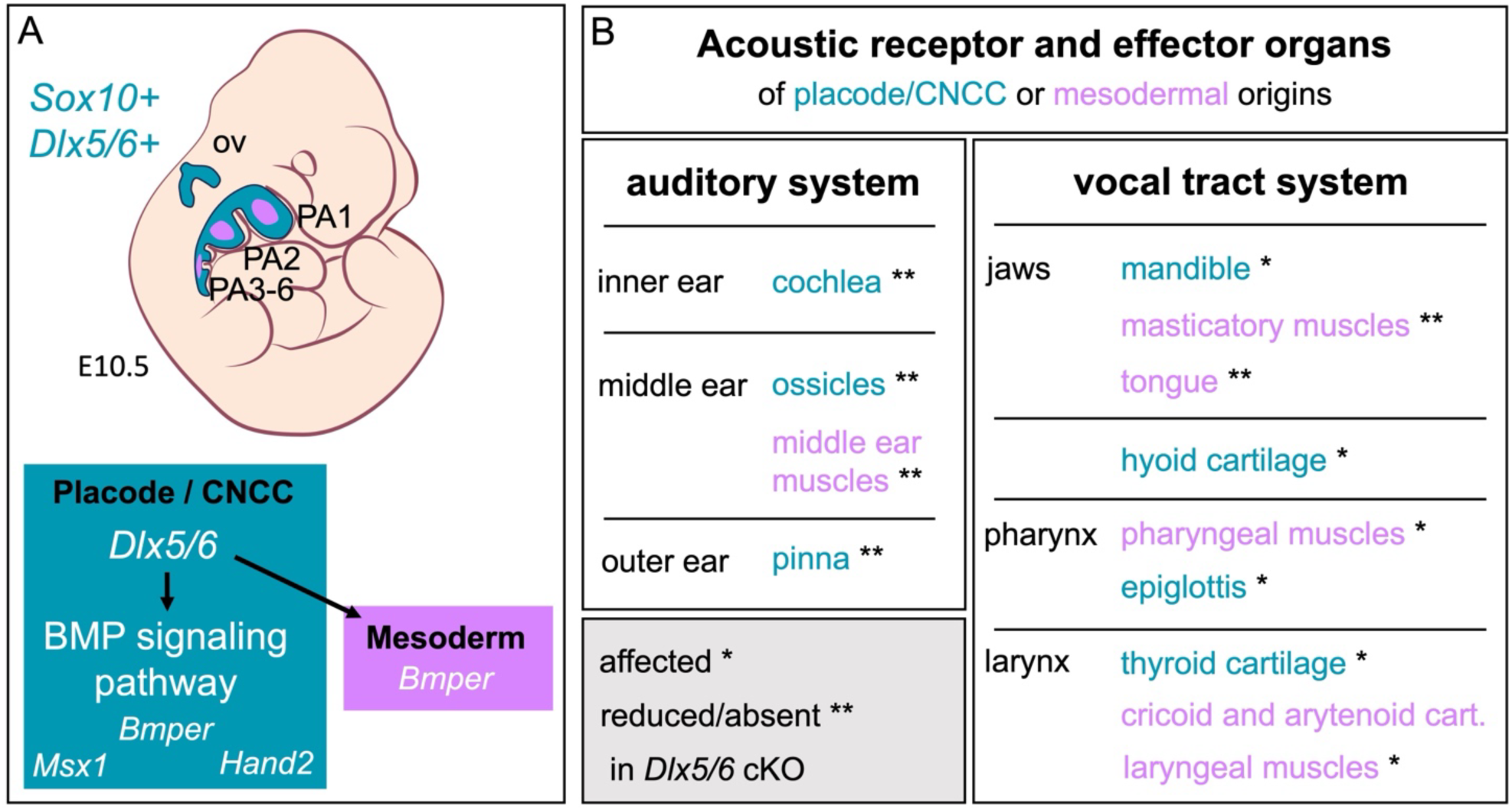
A common Sox10-Dlx5/6-BMP axis acts to coordinate the formation of both auditory and vocal tract systems. (A) Schematic views of *Sox10* and *Dlx5/6* co-expression in the otic vesicle (ov) and pharyngeal arches in E10.5 mouse embryos (in blue), and role of Dlx5/6 in regulating BMP signaling in placode and CNCC derivatives (in blue) and PA mesodermal cores (in pink). (B) Summary of the phenotype resulting from *Dlx5/6* conditional mutation affecting both acoustic receptor and effector organs of placode/CNCC (in blue) or mesodermal origins (in pink).

### Dlx5/6 pleiotropic role may have supported acoustic communication diversification

The *Dlx* gene family has an ancient origin in bilaterians and is present in all vertebrate clades^16^. Expression analyses in chick, xenopus and fish embryos revealed that *Dlx5/6* expression in pharyngeal arches and otic vesicles is conserved across vertebrate species^44–48^. Our data show that *Dlx5/6* genes control simultaneously the formation of all structures composing the vocal tract and auditory systems (Fig. 5), receptor and effector organs for acoustic communication. This fully corresponds to the concept of pleiotropy since a single locus affects independent phenotypic traits^24,25^.

It has been shown that acoustic communication has a single origin in the land vertebrates^1,2^ and that the larynx is the main site of vocal production within the vertebrate family^4^. The mammalian clade comprises diverse ecological niches, a diversity that is explained by the rapid adaptive radiation during “the age of mammals” and terrestrial changes like continental rearrangements^4,49^. Mammals show a wide variety of acoustic transmission varying from infra to ultrasounds, possibly originating from the adaptation to new environments and for predator avoidance^50^. The differences in tone and frequency can be explained by the morphological diversity of the larynx in mammals as well as by the adaptation of the receptor organ capable of hearing infra or ultrasounds^3,4^. It has been suggested that mammalian vocal characteristics and hearing sensitivity co-evolved in the forest mammals, following the sensory drive hypothesis^51^. One of the characteristics of the mammalian clade is the presence of ossicles of CNCC origin. Their morphology is dependent on environment and behavior making the auditory apparatus competent to pick up a wider variety of frequencies^50^. Beyond sound emission and reception, a study revealed that targeted invalidation of *Dlx5/6* in GABAergic neurons results in behavioural alterations including hyper vocalisation and increased socialisation in mice^52^. In humans, *DLX5* was proposed to have played a role in the evolution of the human’s linguistic capacity and skull globularization^53^. We propose that the pleiotropic role of Dlx5/6 would have supported the co-adaptation of vocal and auditory acoustic organs but also cognitive communication capacities in mammals.

Our *Dlx5/6* mutants display malformations that also affect traits of domestication^54^. It has been proposed that the “Neural Crest Domestication Syndrome” (NCDS) results potentially from a reduction of CNCC-derived tissues of behavioural relevance and inducing morphological changes of the jaw, ear or larynx^54–56^. In domesticated mammalian species such as cats and dogs, pets interact with humans with acoustic emissions that are not recorded in the wild. The pleiotropic role of *Dlx5/6* in CNCC and otic placode development may support the morphological changes predicted by the NCDS by adapting the auditory and vocal systems to assure pet-human interactions.

### Incidence for understanding the etiology of human syndromic forms

In humans, the Split Hand/Split Foot malformation 1 (SHFM1, OMIM #183600) is characterized by ectrodactyly caused by pathogenic variants affecting the DLX5/6 locus^57^. *Dlx5/6* mutant mice have been investigated to understand the etiology of the limb phenotype in SHFM1^58^. In patients, the limb malformations can be associated to other defects including cranial malformations, hearing loss and intellectual disabilities (OMIM #220600). It has been proposed that a minimal SHFM1 chromosomal region containing *DLX5/6* enhancers would be compromised and related to associated phenotypes^57^. Our data on the pleiotropic role of *Dlx5/6* in vocal tract and otic development are thus relevant to better understand the hearing loss and craniofacial syndromic features associated with the limb phenotype in SHFM1 patients.

## Acknowledgements

We are grateful to Giovanni Levi and Nicolas Narboux-Nême for discussion and their insightful comments on the manuscript, as well as for the donation of *Dlx5/6^flox/flox^* mice.

We thank the staff of the animal facility, Mr. Stéphane Sosinski and Mr. Fabien Uridat, for their precious and ethical care of the mouse lines.

This work was supported by the National Museum of Natural History’s grants ‘MorphoVox’ and ‘Axonton’, and the Agence Nationale de la Recherche’s grant ‘MorphoNeck’ (ANR-21-CE13-0025) (EH, FSG). The CzechNanoLab Research Infrastructure was supported by MEYS CR (LM2023051) (MK, TZ, JK).

## Author contributions

FSG: Investigation, data acquisition, visualisation, funding acquisition, writing original draft, writing review and editing

VB: Investigation, data acquisition, validation, visualisation

CC: Data acquisition, visualisation

AF: Animal colony management

MK: data acquisition, visualisation

TZ & JK: Resources

EH: Conceptualization, methodology, investigation, visualisation, validation, project administration, funding acquisition, writing original draft, writing review and editing

## Materials and Methods

### Animals

Procedures involving animals were conducted in accordance with European Community (Council Directive 86/609) and French Agriculture Ministry directives (Council Directive 87– 848). The project was approved by the “Cuvier” ethical committee of the french National Museum of National History (MNHN) and validated by the French Ministry of Agriculture (approval APAFIS#26087-2020061614327140).

All animals were housed in light, temperature (21°C) and humidity (50%-60%) controlled conditions. Food and water were available *ad libitum*. Mice were back-crossed and maintained with a B6D2F1/J background. All genotyping was made by polymerase chain reaction (PCR) on ear biopsies.

Males carrying *loxP* sequences flanking the exon 2-containing homeobox domain of *Dlx5* and *Dlx6* genes (*Dlx5/6^flox/flox^*)^29^ were crossed with *Sox10^Cre/+^* females^28^ to invalidate *Dlx5/6* (*Sox10^Cre/+^ : Dlx5/6^flox/flox^*). For the genetic lineage tracing, *Sox10^Cre/+^ : Dlx5/6^flox/+^* females were crossed with males carrying *Dlx5/6^flox/flox^* and the *Rosa^lox-stop-lox-lacZ^* reporter^30^ (*Dlx5/6^flox/flox^ : Rosa^lacZ/lacZ^*).

For *Dlx5* expression analysis, males carrying the lacZ reporter (*Dlx5^lacZ/+^*)^18^ were crossed with B6D2F1/J females.

Samples at E10.5, E11.5 and E17.5 stages were fixed in 4% paraformaldehyde (PFA) diluted in PBS overnight at 4°C and washed in PBS. Foetuses at stage E14.5 and E16.5 were fixed for 3 hours in 4% PFA diluted in PBS and 0.5% Triton (PBSTr) at 4°C and washed twice in Tween 0.1% diluted in PBS (PBST) and then again overnight at 4°C. For all samples, noon of the day of the vaginal plug was considered as E0.5.

### X-Gal and Immunofluorescence stainings

Whole-mount samples at E9.5, E10.5, E11.5 and E14.5 were collected and fixed in PFA 4% for 3 hours, washed and treated with X-gal to reveal β-galactosidase activity as previously described^7^. For whole-mount immunofluorescence stainings, embryos at E11.5 were collected, fixed and dehydrated in a graded series of methanol (MeOH). They were then permeabilized in Dent’s solution (80% MeOH – 20% DMSO) overnight at 4°C and rehydrated in graded MeOH series and washed in PBS. The non-specific antigenic sites were blocked using a blocking solution (BS, PBS / 20% goat serum / 3% BSA / 0.5 % Triton) for 3 hours at room temperature (RT). The embryos were incubated with primary antibody diluted in BS for 2-3 days with gentle shaking at 4°C, washed several times over day in PBST and incubated with secondary antibodies also diluted in BS for 2-3 days with gentle shaking at 4°C in the dark. They were finally washed several times over day and cleared following the Cubic protocol as previously described^59^.

For immunofluorescence stainings on sections, foetuses were cryopreserved in 30% sucrose PBS and embedded in OCT for 20μm sectioning using a Leica cryostat. Cryosections were dried for 30 minutes, washed in PBS and immersed in BS for 2 hours at RT. Primary antibodies were diluted in BS and sections were incubated overnight at 4°C. The sections were then washed and incubated with secondary antibodies diluted in BS for 2 hours at RT in the dark. Finally, the sections were washed in PBS and mounted with Fluoromount (Fisher Scientific, 15586276) for analysis.

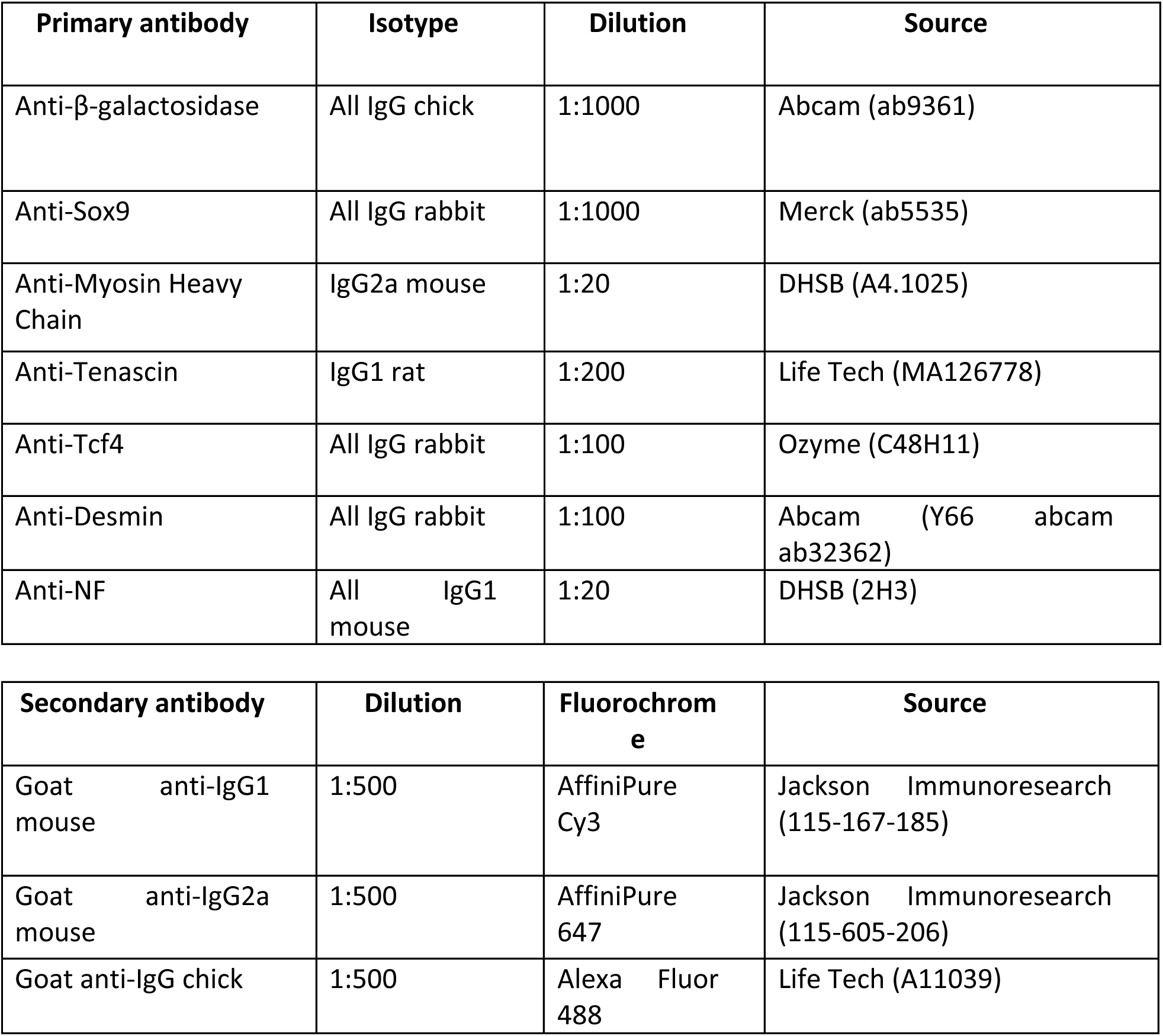
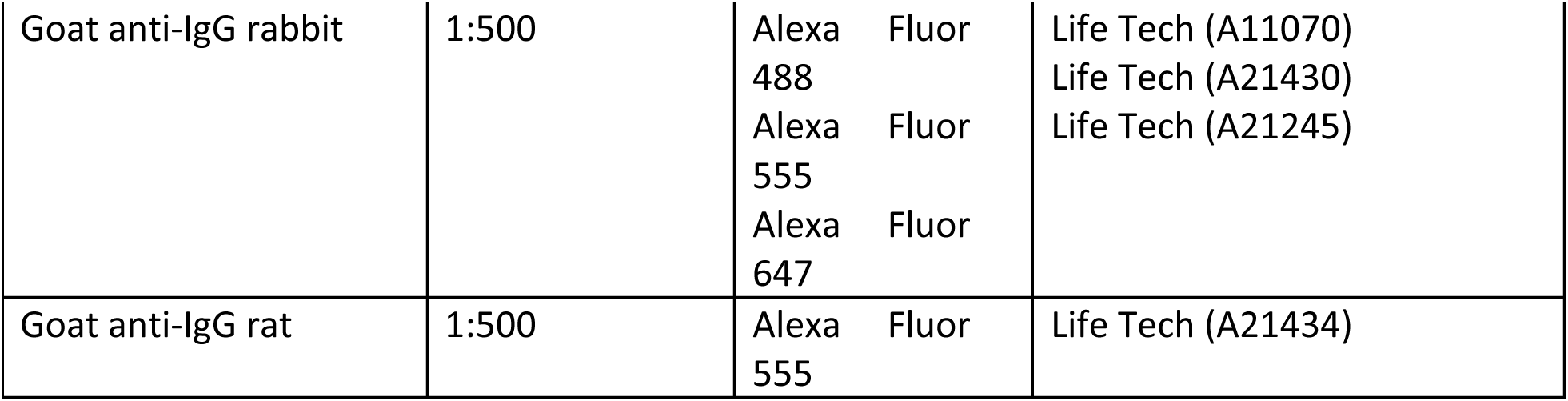

### Immunofluorescence acquisitions and analysis

Whole mount samples for lightsheet imaging were prepared by CUBIC L-R(N) clearing protocol^59^, after fixation and immunofluorescence staining. Samples were included in agarose low melting temperature 2 % (A9414, Sigma-Aldrich). We first incubated samples in 10 mL of 50% CUBIC-L reagent composed of 10% (w/w) N-butyldiethanolamine (471240, Sigma-Aldrich) and 10% (w/w) triton X-100 (Sigma-Aldrich) diluted in MilliQ water for 2 days at RT, for 2 days in 10 mL of 100% CUBIC-L at RT, and washed 3 times overnight in 10 mL of PBS. They were then incubated in 10 mL of 50% CUBIC-R(N) reagent composed of 45% (w/w) Antipyrine (A5882, Sigma-Aldrich), 30% (w/w) Nicotinamide (A15970, Thermo Fisher) an 0,5% (w/w) N-butyldiethanolamine (471240, Sigma-Aldrich) diluted in MilliQ water for 2 days at RT and for 2 days in 10 mL of 100% CUBIC-L at RT. For lightsheet imaging, the samples were incubated in a fresh CUBIC-R(N) reagent in the tank and incubated for a few minutes for refractive index homogenization.

Lightsheet imaging was performed with the Alpha3 system (PhaseView, France), equipped with a 10× XL Plan N (XLPLN10XSVMP, Olympus, Japan) clearing objective with refractive index adaptive collar (RI 1.33–1.52), a sCMOs Orca Flash4 camera (Hamamatsu, Japan). 2.20 and the acquisition software QtSPIM. For 561 nm excitation laser, the ET600/50m emission filter was used and for the 638 nm excitation laser, the emission filter ET670/50m.

### Image reconstruction and analysis

Confocal and macroscope images were reconstructed and analyzed using Fiji ImageJ (Java version 1.8.0) and ZEN Blue 3.5 (Zeiss). Airyscan and lightsheet images were converted in arivis SIS Converter 3.1.1 and analyzed with arivis Vision4D 3.0.1 software (arivis AG, Germany).

| System | Type | Objective (NA) | Lasers |
| --- | --- | --- | --- |
| LSM 980 –<br>AiryScan 2<br>detector Zeiss | Confocal –<br>AiryScan | EC-Plan Neofluar 10X dry (0.3 NA) or Plan-Apochromat water 20X (0.8 NA) or 63X C-Apochromat oil (1.2 NA) | 405 – 488 – 561 – 639 nm |
| Macro-Apotome<br>AxioZoom V16<br>Zeiss | Macroscope | PlanNeoFluar Z 1.0X oil (0.25 NA) or PlanNeoFluar Z 2,3X (0.57 NA) | Excitation by fluorescent lamp with adapted filter block (DAPI, GFP, DsRed, Cy5) |
| Lightsheet<br>Alpha3<br>PhaseView | Lightsheet | XLPLN 10X Olympus objective (0.6 NA) multi-immersion refractive index adaptive collar (RI 1.33 – 1.52) | 405 – 488 – 561 – 635 nm |

### X-ray computed microtomography (micro-CT) acquisitions and analysis

E17.5 foetuses were fixed overnight in PFA 4% and washed in several baths of PBST over day. They were dehydrated in upgrading EtOH and MeOH series and soaked in Phosphotungstic acid (PTA) 1.5% diluted in MeOH 90% for 4 days. The PTA solution was changed everyday with a fresh solution to ensure optimal penetration of the contrast agent. The micro-CT scanning was done by X-ray micro-CT system GE Phoenix v|tome|x L 240 (Waygate Technologies / Baker Hughes Digital Solutions GmbH, Wunstorf) equipped with a 180 kV/15 W maximum power nanofocus X-ray tube and a high-contrast flat panel dynamic detector 41|100 with 4000 × 4000 pixels and a pixel size of 100 × 100 μm. The exposure time was 800 ms in 2000 positions over 360°. Three projections were captured in each position and an average of the signal was used to improve the signal-to-noise ratio. The microCT scan was carried out at 600 kV acceleration voltage and with 200 μA X-ray tube current. The beam was filtered by a 0.2 mm-thick aluminium filter to avoid beam hardening artefacts. The isotropic voxel size of obtained volumes was 9 μm for all four samples: 2 control and 2 *Dlx5/6* cKO. The tomographic reconstruction was performed using GE phoenix datos|x 2.0 software (Waygate Technologies / Baker Hughes Digital Solutions GmbH, Wunstorf, Germany). Reconstructed slices were imported to software Avizo 7.1 (Thermo Fisher Scientific, Waltham, MA, USA) for semi-automatic segmentation as previously described in our previous work^60^. The muscles and other anatomical structures of interest were outlined by the operator in every 3rd to 5th slice depending on the complexity of the structure and the rest was calculated by linear interpolation between manually outlined slices. The segmented structures were then transferred to polygonal mesh (STL format) and imported to VG Studio MAX 3.5 (Volume Graphics GmbH, Heidelberg, Germany) for further visualisation and analysis.

### Preparation of cDNA and RNA probes from embryos

RNA extractions were done using PureLink RNA Mini Kit with Trizol following the kit guidelines (ThermoFisher, 12183018A) and turned into cDNA libraries following the SuperScript III First-Strand Synthesis SuperMix for qRT-PCR (ThermoFisher, 11752-050) protocol. RNA probes were designed on primer BLAST (https://www.ncbi.nlm.nih.gov/tools/primer-blast/) and synthesised by PCR, following the guidelines of Gel Extraction Kit (Qiagen, 28704), PCR Purification Kit (Qiagen, 28104) and using T7 to generate RNA probes for *in situ* hybridization.

### In situ hybridization

Embryos at E10.5 were collected, fixed, dehydrated in a graded series of MeOH and stored at - 20°C. They were rehydrated progressively in PBST, permeabilized by Proteinase K (10 µg/ml in PBST) and post fixed in PFA 4%. Samples were equilibrated in a solution of 50% Hybridization solution (Hb) - 50% PBST 10 minutes at room temperature under agitation and in 100% Hb for an hour at 70°C. Embryos were then incubated in Hb with 1ng/uL RNA probe overnight at 70°C. After 4 successive washes of 30 minutes with Wash Buffer (WB) at 70°C, they were equilibrated in a graded series of TBST at room temperature. Embryos were blocked with Blocking Buffer (BB) and incubated overnight at 4°C in Antibody Solution (AbS) containing anti-DIG antibodies coupled with alkaline phosphatase (Merk, 11093274910). Samples were washed in TBST several times followed by 3 NTMT washes and revealed with BM-Purple (Merk, 11442074001) until they reached proper staining contrasts. The Hb, WB, TBST, BB, SAc and NTMN composition details are presented in Supplementary data 1.

---

**Supp. Fig. 1.**
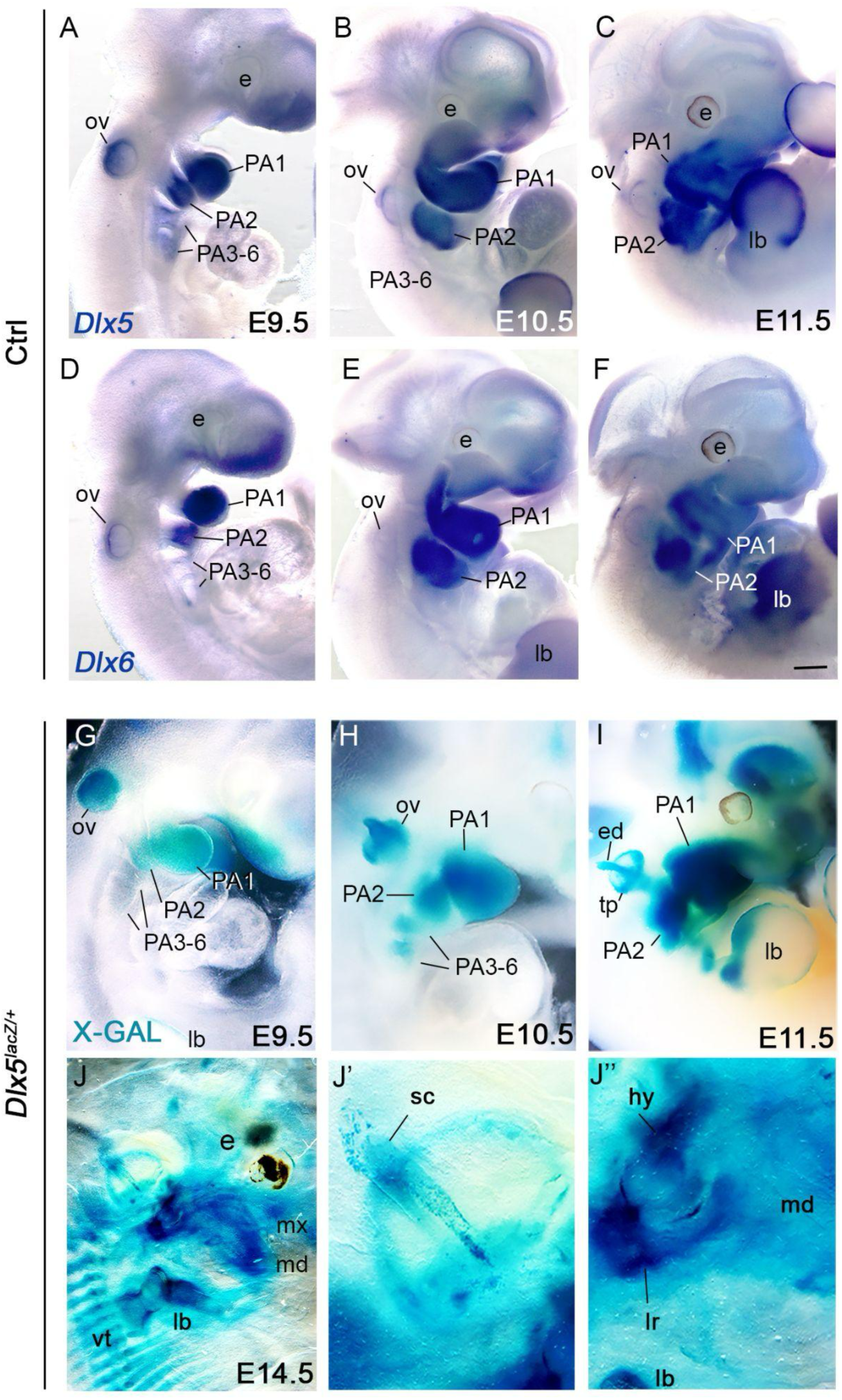
*Dlx5* and *Dlx6* expressions during mouse development. (A-F) *In situ* hybridization for *Dlx5* and *Dlx6* at E9.5, E10.5 and E11.5 in control embryos. Note that the genes show similar spatio-temporal expression profiles within PA CNCC and otic vesicle. (G-J) *Dlx5-lacZ* reporter expression in E9.5, E10.5, E11.5 embryos overlap *Dlx5* and *Dlx6* expression profile but better highlights expression in posterior PAs (PA3-6). In clarified E14.5 *Dlx5^lacZ/+^* fetuses, β-gal expression is activated in the developing skeleton. Magnifications of the ear (J’) and laryngeal regions (J’’). Abbreviations: e, eye; lb, limb bones; mx, maxilar; mb, mandibular; vt, vertebrae; sc, semicircular canals; hy, hyoid cartilage; lr, laryngeal cartilage; PA, pharyngeal arch; OV, otic vesicle. Scale bar in F for A, D, G 150 µm, for B, E, H 200 µm, for C, F, I 300 µm, for J 500 µm, for J’-J” 150 µm.

**Supp. Fig. 2.**
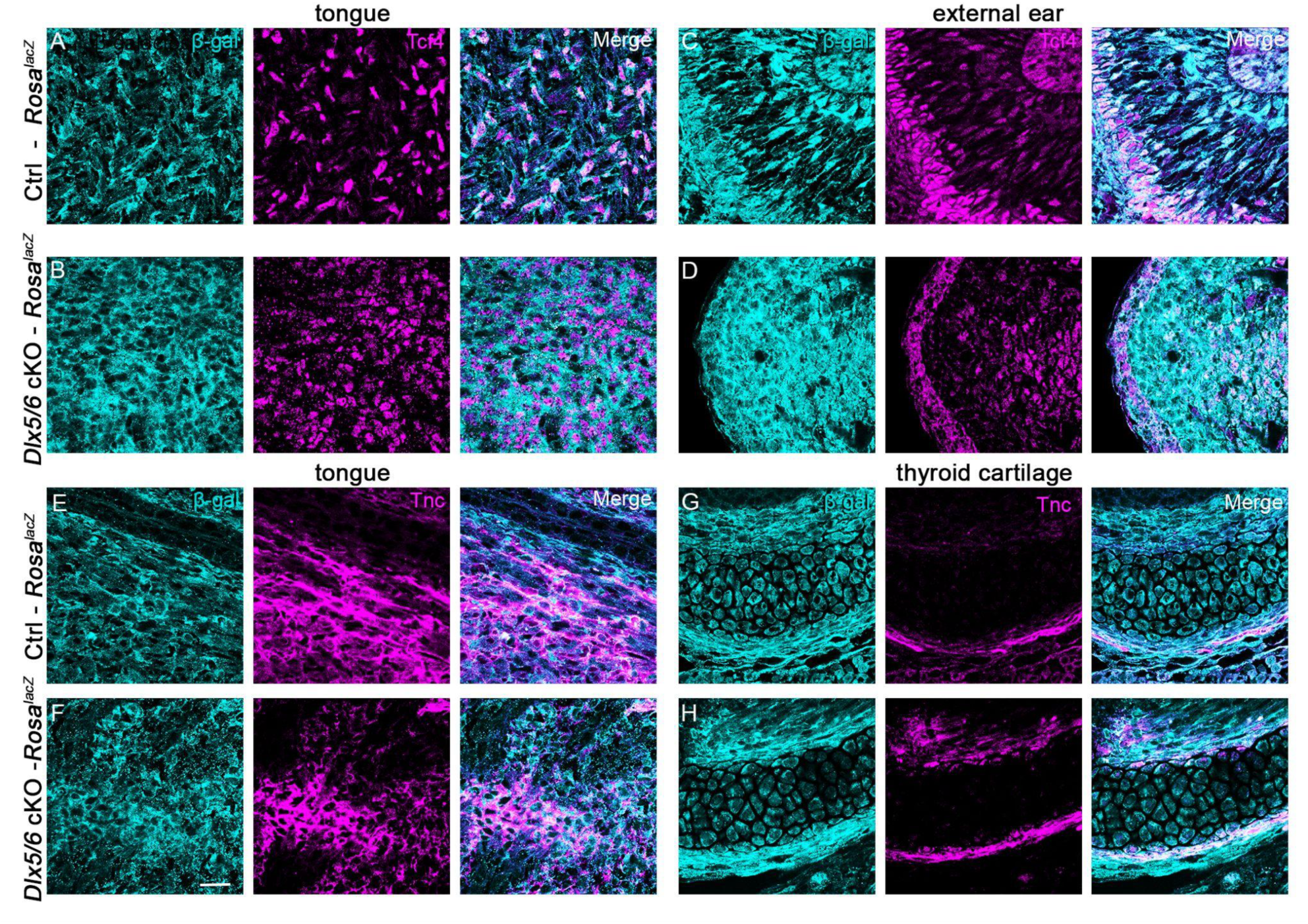
Invalidation of *Dlx5/6* affects CNCC patterning but not CNCC fate. (A-H) Immunofluorescence stainings for β-gal and for Tcf4 and Tnc, markers of CNCC-derived fibroblasts and tendons at the tongue, external ear and thyroid levels in control and *Dlx5/6* mutant conditions. (A-B, E-F). In the tongue, β-gal-positive cells show defects of patterning but keep their fibroblast and tendinous identities (C-D), as they do at external ear and thyroid levels (C-D, E-H) (n=3). Scale bar in F for A-H 20 µm.

## Supplementary data

**Supplementary data 1.** Compositions of solutions used for *in situ* hybridization

**Supplementary data 2-3**. Interactive 3D PDF of the 3D reconstructions of control and *Dlx5/6* mutant phenotypes of vocal tract and auditory systems to complete data shown in Figure 1 E-H’.

**Supplementary videos 1-2**. Interactive movies of the neuromuscular system of control and *Dlx5/6* mutant embryos to complete data shown in Figure 3.

**Supplementary data 1.**
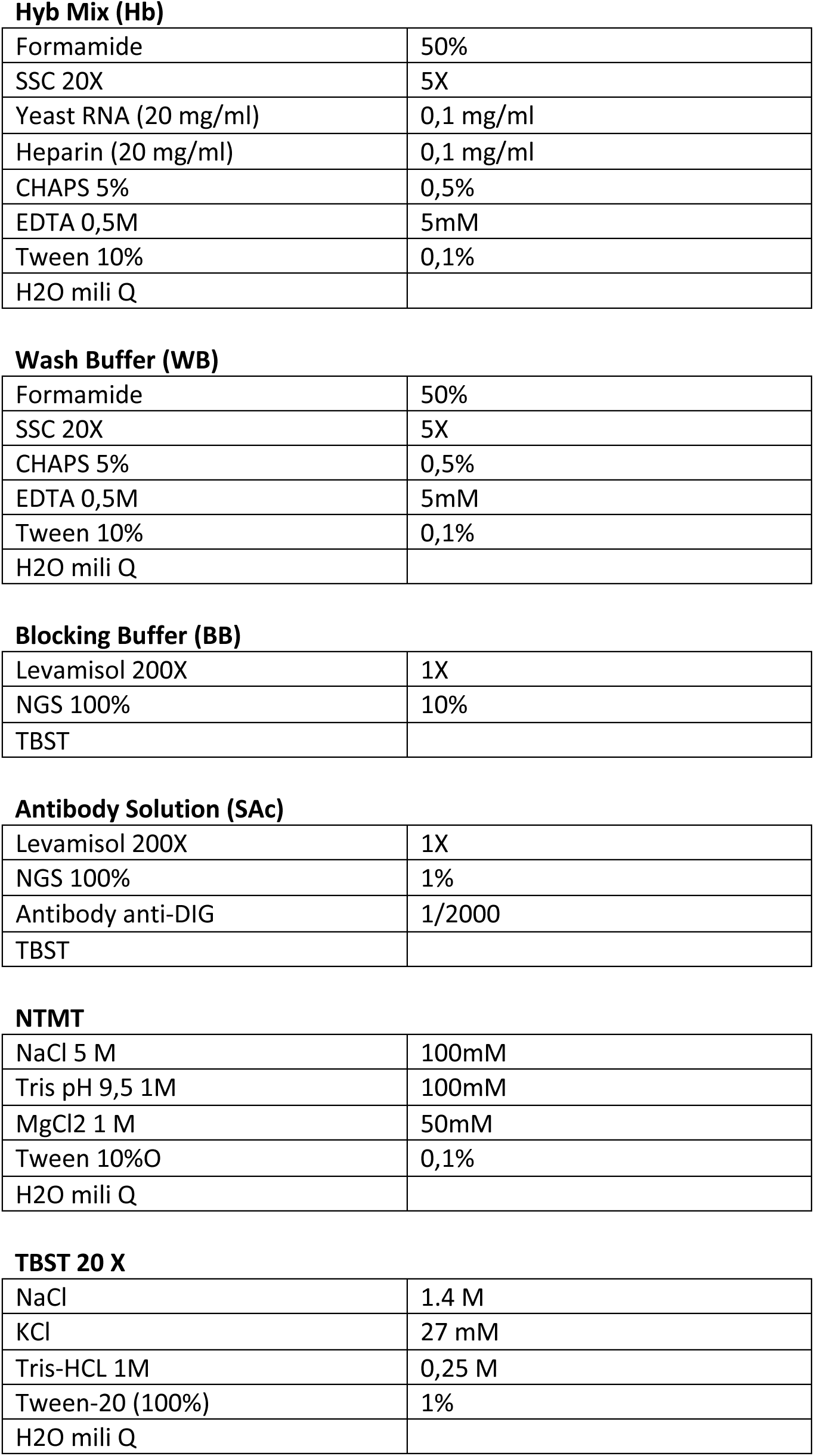
Compositions of solutions used for *in situ* hybridization.

|  |  |
| --- | --- |
| Formamide | 50% |
| SSC 20X | 5X |
| Yeast RNA (20 mg/ml) | 0,1 mg/ml |
| Heparin (20 mg/ml) | 0,1 mg/ml |
| CHAPS 5% | 0,5% |
| EDTA 0,5M | 5mM |
| Tween 10% | 0,1% |
| H2O mili Q |  |
| **Wash Buffer (WB)** |  |
| Formamide | 50% |
| SSC 20X | 5X |
| CHAPS 5% | 0,5% |
| EDTA 0,5M | 5mM |
| Tween 10% | 0,1% |
| H2O mili Q |  |
| **Blocking Buffer (BB)** |  |
| Levamisol 200X | 1X |
| NGS 100% | 10% |
| TBST |  |
| **Antibody Solution (SAc)** |  |
| Levamisol 200X | 1X |
| NGS 100% | 1% |
| Antibody anti-DIG | 1/2000 |
| TBST |  |
| **NTMT** |  |
| NaCl 5 M | 100mM |
| Tris pH 9,5 1M | 100mM |
| MgCl2 1 M | 50mM |
| Tween 10%O | 0,1% |
| H2O mili Q |  |
| **TBST 20 X** |  |
| NaCl | 1.4 M |
| KCl | 27 mM |
| Tris-HCL 1M | 0,25 M |
| Tween-20 (100%) | 1% |
| H2O mili Q |  |

## References

1 Chen, Z. & Wiens, J. J. The origins of acoustic communication in vertebrates. Nat Commun 11, 369 (2020). 10.1038/s41467-020-14356-3

2 Jorgewich-Cohen, G. et al. Common evolutionary origin of acoustic communication in choanate vertebrates. Nat Commun 13, 6089 (2022). 10.1038/s41467-022-33741-8

3 Ladich, F. & Winkler, H. Acoustic communication in terrestrial and aquatic vertebrates. J Exp Biol 220, 2306–2317 (2017). 10.1242/jeb.132944

4 Taylor, A. M., Charlton, B. D. & Reby, D. in Vertebrate Sound Production and Acoustic Communication (eds Roderick A. Suthers, W. Tecumseh Fitch, Richard R. Fay, & Arthur N. Popper) 229–259 (Springer International Publishing, 2016).

5 Wilkins, M. R., Seddon, N. & Safran, R. J. Evolutionary divergence in acoustic signals: causes and consequences. Trends Ecol Evol 28, 156–166 (2013). 10.1016/j.tree.2012.10.002

6 Frisdal, A. & Trainor, P. A. Development and evolution of the pharyngeal apparatus. Wiley Interdiscip Rev Dev Biol 3, 403–418 (2014). 10.1002/wdev.147

7 Heude, E. et al. Unique morphogenetic signatures define mammalian neck muscles and associated connective tissues. Elife 7 (2018). 10.7554/eLife.40179

8 Lungova, V. & Thibeault, S. L. Mechanisms of larynx and vocal fold development and pathogenesis. Cell Mol Life Sci 77, 3781–3795 (2020). 10.1007/s00018-020-03506-x

9 Shimizu, M. et al. Probing the origin of matching functional jaws: roles of Dlx5/6 in cranial neural crest cells. Sci Rep 8, 14975 (2018). 10.1038/s41598-018-33207-2

10 Ziermann, J. M., Diogo, R. & Noden, D. M. Neural crest and the patterning of vertebrate craniofacial muscles. Genesis 56, e23097 (2018). 10.1002/dvg.23097

11 Adachi, N., Bilio, M., Baldini, A. & Kelly, R. G. Cardiopharyngeal mesoderm origins of musculoskeletal and connective tissues in the mammalian pharynx. Development 147 (2020). 10.1242/dev.185256

12 Tabler, J. M. et al. Cilia-mediated Hedgehog signaling controls form and function in the mammalian larynx. Elife 6 (2017). 10.7554/eLife.19153

13 Anthwal, N. & Thompson, H. The development of the mammalian outer and middle ear. J Anat 228, 217–232 (2016). 10.1111/joa.12344

14 Fuchs, J. C. & Tucker, A. S. in Current Topics in Developmental Biology Vol. 115 (ed Yang Chai) 213–232 (Academic Press, 2015).

15 Heude, E. et al. Jaw muscularization requires Dlx expression by cranial neural crest cells. Proc Natl Acad Sci U S A 107, 11441–11446 (2010). 10.1073/pnas.1001582107

16 Neidert, A. H., Virupannavar, V., Hooker, G. W. & Langeland, J. A. Lamprey Dlx genes and early vertebrate evolution. Proc Natl Acad Sci U S A 98, 1665–1670 (2001). 10.1073/pnas.98.4.1665

17 Panganiban, G. & Rubenstein, J. L. Developmental functions of the Distal-less/Dlx homeobox genes. Development 129, 4371–4386 (2002). 10.1242/dev.129.19.4371

18 Acampora, D. et al. Craniofacial, vestibular and bone defects in mice lacking the Distal-less-related gene Dlx5. Development 126, 3795–3809 (1999).

19 Depew, M. J., Lufkin, T. & Rubenstein, J. L. Specification of jaw subdivisions by Dlx genes. Science 298, 381–385 (2002).

20 Beverdam, A. et al. Jaw transformation with gain of symmetry after Dlx5/Dlx6 inactivation: mirror of the past? Genesis 34, 221–227 (2002).

21 Narboux-Neme, N., Ekker, M., Levi, G. & Heude, E. Posterior axis formation requires Dlx5/Dlx6 expression at the neural plate border. PLoS One 14, e0214063 (2019). 10.1371/journal.pone.0214063

22 Merlo, G. R. et al. The Dlx5 homeobox gene is essential for vestibular morphogenesis in the mouse embryo through a BMP4-mediated pathway. Dev Biol 248, 157–169 (2002).

23 Robledo, R. F. & Lufkin, T. Dlx5 and Dlx6 homeobox genes are required for specification of the mammalian vestibular apparatus. Genesis 44, 425–437 (2006). 10.1002/dvg.20233

24 Stearns, F. W. One hundred years of pleiotropy: a retrospective. Genetics 186, 767–773 (2010). 10.1534/genetics.110.122549

25 Wang, Z., Liao, B. Y. & Zhang, J. Genomic patterns of pleiotropy and the evolution of complexity. Proc Natl Acad Sci U S A 107, 18034–18039 (2010). 10.1073/pnas.1004666107

26 Zhang, J. Patterns and evolutionary consequences of pleiotropy. Annu Rev Ecol Evol Syst 54, 1–19 (2023). 10.1146/annurev-ecolsys-022323-083451

27 Betancur, P., Sauka-Spengler, T. & Bronner, M. A Sox10 enhancer element common to the otic placode and neural crest is activated by tissue-specific paralogs. Development 138, 3689–3698 (2011). 10.1242/dev.057836

28 Jacques-Fricke, B. T., Roffers-Agarwal, J. & Gammill, L. S. DNA methyltransferase 3b is dispensable for mouse neural crest development. PLoS One 7, e47794 (2012). 10.1371/journal.pone.0047794

29 de Lombares, C. et al. Dlx5 and Dlx6 expression in GABAergic neurons controls behavior, metabolism, healthy aging and lifespan. Aging (Albany NY) 11, 6638–6656 (2019). 10.18632/aging.102141

30 Friedrich, G. & Soriano, P. Promoter traps in embryonic stem cells: a genetic screen to identify and mutate developmental genes in mice. Genes Dev 5, 1513–1523 (1991). 10.1101/gad.5.9.1513

31 Sugimoto, T. et al. Three-Dimensional Visualization of Developing Neurovascular Architecture in the Craniofacial Region of Embryonic Mice. Anat Rec (Hoboken) 298, 1824–1835 (2015). 10.1002/ar.23179

32 Vermeiren, S., Bellefroid, E. J. & Desiderio, S. Vertebrate Sensory Ganglia: Common and Divergent Features of the Transcriptional Programs Generating Their Functional Specialization. Front Cell Dev Biol 8, 587699 (2020). 10.3389/fcell.2020.587699

33 Sajan, S. A., Rubenstein, J. L., Warchol, M. E. & Lovett, M. Identification of direct downstream targets of Dlx5 during early inner ear development. Hum Mol Genet 20, 1262–1273 (2011). 10.1093/hmg/ddq567

34 Jeong, J. et al. Dlx genes pattern mammalian jaw primordium by regulating both lower jaw-specific and upper jaw-specific genetic programs. Development 135, 2905–2916 (2008).

35 Coles, E., Christiansen, J., Economou, A., Bronner-Fraser, M. & Wilkinson, D. G. A vertebrate crossveinless 2 homologue modulates BMP activity and neural crest cell migration. Development 131, 5309–5317 (2004). 10.1242/dev.01419

36 Ikeya, M. et al. Essential pro-Bmp roles of crossveinless 2 in mouse organogenesis. Development 133, 4463–4473 (2006). 10.1242/dev.02647

37 Moser, M. et al. BMPER, a novel endothelial cell precursor-derived protein, antagonizes bone morphogenetic protein signaling and endothelial cell differentiation. Mol Cell Biol 23, 5664–5679 (2003). 10.1128/MCB.23.16.5664-5679.2003

38 Charite, J. et al. Role of Dlx6 in regulation of an endothelin-1-dependent, dHAND branchial arch enhancer. Genes Dev 15, 3039–3049 (2001).

39 Levi, G. et al. Msx1 and Dlx5 act independently in development of craniofacial skeleton, but converge on the regulation of Bmp signaling in palate formation. Mech Dev 123, 3–16 (2006). 10.1016/j.mod.2005.10.007

40 Funato, N., Kokubo, H., Nakamura, M., Yanagisawa, H. & Saga, Y. Specification of jaw identity by the Hand2 transcription factor. Sci Rep 6, 28405 (2016). 10.1038/srep28405

41 Kuratani, S. et al. Middle ear defects associated with the double knock out mutation of murine goosecoid and Msx1 genes. Cell Mol Biol (Noisy-le-grand) 45, 589–599 (1999).

42 Wang, R. N. et al. Bone Morphogenetic Protein (BMP) signaling in development and human diseases. Genes & Diseases 1, 87–105 (2014). 10.1016/j.gendis.2014.07.005

43 Vincentz, J. W. et al. Exclusion of Dlx5/6 expression from the distal-most mandibular arches enables BMP-mediated specification of the distal cap. Proc Natl Acad Sci U S A 113, 7563–7568 (2016). 10.1073/pnas.1603930113

44 Ellies, D. L. et al. Relationship between the genomic organization and the overlapping embryonic expression patterns of the zebrafish dlx genes. Genomics 45, 580–590 (1997). 10.1006/geno.1997.4978

45 Keer, S. et al. Bop1 is required to establish precursor domains of craniofacial tissues. Genesis 62, e23580 (2024). 10.1002/dvg.23580

46 Papalopulu, N. & Kintner, C. Xenopus Distal-less related homeobox genes are expressed in the developing forebrain and are induced by planar signals. Development 117, 961–975 (1993). 10.1242/dev.117.3.961

47 Sohail, A. & Bendall, A. J. DLX gene expression in the developing chick pharyngeal arches and relationship to endothelin signaling and avian jaw patterning. Dev Dyn 253, 255–271 (2024). 10.1002/dvdy.653

48 Talbot, J. C., Johnson, S. L. & Kimmel, C. B. hand2 and Dlx genes specify dorsal, intermediate and ventral domains within zebrafish pharyngeal arches. Development 137, 2507–2517 (2010). 10.1242/dev.049700

49 Meredith, R. W. et al. Impacts of the Cretaceous Terrestrial Revolution and KPg extinction on mammal diversification. Science 334, 521–524 (2011). 10.1126/science.1211028

50 Le Maître, A., Grunstra, N. D. S., Pfaff, C. & Mitteroecker, P. Evolution of the Mammalian Ear: An Evolvability Hypothesis. Evol Biol 47, 187–192 (2020). 10.1007/s11692-020-09502-0

51 Charlton, B. D., Owen, M. A. & Swaisgood, R. R. Coevolution of vocal signal characteristics and hearing sensitivity in forest mammals. Nat Commun 10, 2778 (2019). 10.1038/s41467-019-10768-y

52 Levi, G. et al. DLX5/6 GABAergic Expression Affects Social Vocalization: Implications for Human Evolution. Mol Biol Evol 38, 4748–4764 (2021). 10.1093/molbev/msab181

53 Boeckx, C. & Benítez-Burraco, A. The shape of the human language-ready brain. Front Psychol 5, 282 (2014). 10.3389/fpsyg.2014.00282

54 Wilkins, A. S., Wrangham, R. W. & Fitch, W. T. The "domestication syndrome" in mammals: a unified explanation based on neural crest cell behavior and genetics. Genetics 197, 795–808 (2014). 10.1534/genetics.114.165423

55 Lesch, R. & Fitch, W. T. The domestication of the larynx: The neural crest connection. J Exp Zool B Mol Dev Evol 342, 342–349 (2024). 10.1002/jez.b.23251

56 Wilkins, A. S., Wrangham, R. & Fitch, W. T. The neural crest/domestication syndrome hypothesis, explained: reply to Johnsson, Henriksen, and Wright. Genetics 219 (2021). 10.1093/genetics/iyab098

57 Birnbaum, R. Y. et al. Functional characterization of tissue-specific enhancers in the DLX5/6 locus. Hum Mol Genet 21, 4930–4938 (2012). 10.1093/hmg/dds336

58 Robledo, R. F., Rajan, L., Li, X. & Lufkin, T. The Dlx5 and Dlx6 homeobox genes are essential for craniofacial, axial, and appendicular skeletal development. Genes Dev 16, 1089–1101 (2002). 10.1101/gad.988402

59 Matsumoto, K. et al. Advanced CUBIC tissue clearing for whole-organ cell profiling. Nat Protoc 14, 3506–3537 (2019). 10.1038/s41596-019-0240-9

60 Tesarova, M. et al. An interactive and intuitive visualisation method for X-ray computed tomography data of biological samples in 3D Portable Document Format. Sci Rep 9, 14896 (2019). 10.1038/s41598-019-51180-2

